# Noradrenergic alpha-2A receptor activation suppresses courtship vocalization in male Japanese quail

**DOI:** 10.1101/2021.04.01.438152

**Authors:** Yasuko Tobari, Ami Masuzawa, Norika Harada, Kenta Suzuki, Simone L. Meddle

## Abstract

Male Japanese quail produce high-frequency crow vocalizations to attract females during the breeding season. The nucleus of intercollicularis (ICo) is the midbrain vocal center in birds and electrical stimulation of the ICo produces calls that include crowing. Noradrenaline plays a significant role in sexual behavior but the contribution of noradrenaline in the control of courtship vocalizations in quail has not been well established. Using dose-dependent intracerebroventricular injection of clonidine, an α2-adrenergic receptor- specific agonist, crowing vocalization was immediately suppressed. At the same time as crow suppression by clonidine there was a reduction of immediate early gene, zenk mRNA, in the ICo; no zenk mRNA expression was detected in the dorsomedial division of the nucleus. Using histochemistry, we determined that the ICo receives noradrenergic innervation and expresses α2A-adrenergic receptor mRNA. Taken together, these data suggest that noradrenaline regulates courtship vocalization in quail, possibly via the alpha2A- adrenergic receptor expressed on ICo neurons.

## 1. Introduction

Vocal behavior is commonly used by vertebrates for intraspecific communication. Vocalizations are emitted according to the behavioral context, and under some circumstances, vocalization abruptly stops in respond to a change in situation. Crowing vocalization is one of the most characteristic courtship behaviors exhibited by male Japanese quail (*Coturnix japonica*), and data from the field suggest that males crow primarily in the absence of females [1]. In captivity, crowing, which can reach 95 dB, consists of two to three syllables [2] and is performed by sexually mature, individually housed males that are at least 32 days old [3]. Playback experiments have shown that females, but not males, exhibit positive phonotaxis to crowing [2]. Crowing behavior is context dependent as when males are in the presence of a female, they instantly stop crowing and actively approach her for copulation [4-7] suggesting that crowing functions to attract females. The presence of a female tends to be sufficient to suppress male crowing in Japanese quail, but to date the neurochemical pathway involved in the suppression of crowing in the brain has not been identified.

The mesencephalic nucleus intercollicularis (ICo) is a crucial component of the vocal control system in birds including Galliforme species [8]. Electrical stimulation of the ICo alone produces a type of calling with some characteristic acoustic features of crowing [9-11]. Perturbation of the neural activity of the ICo results in disruption of ongoing crowing behavior in sexually mature male quail [12] and vocal activity is reduced or even eliminated following bilateral lesions of the ICo in the quail [13] and ring dove (*Streptopelia risoria*, [14]. Receptor binding experiments in quail show that there is a high density of α2-adrenergic receptors in the ICo [15, 16], but a low concentration of α1-adrenergic receptors [17]. Noradrenaline binds to the α2-adrenergic receptor, which is primarily postsynaptic in the ICo of quail [18]. The existence of sexual dimorphism in the α2-adrenergic receptor density in the quail ICo suggests that noradrenaline is a possible neurochemical factor underlying the sexually dimorphic vocal behavior. The effect of noradrenaline on call vocalizations has previously been investigated in the ring dove and chicken (*Gallus gallu*s) where pharmacological treatments point to an inhibitory action of noradrenaline on calling behavior [19, 20].

In the current study, we tested the hypothesis that brain noradrenaline acts through α2-adrenergic receptors in the ICo to attenuate crowing in adult male quail in breeding condition. Male quail were injected centrally with noradrenaline, clonidine (α2-adrenergic receptor agonist), or vehicle, and the number of crowing vocalizations in a 1 h period immediately following the injections was quantified. We also tested the hypothesis that crowing suppression is an active inhibitory process within the ICo by quantifying immediate early gene (IEG) expression, which is used as a marker of neuronal activation [21]. The expression of zenk (an acronym of *zif-268, egr-1, ngfI-a*, and *krox-24*) IEG mRNA in the ICo of male quail centrally injected with clonidine or vehicle was quantified. Finally, to test the hypothesis that noradrenaline directly controls ICo neurons, noradrenergic innervation and expression of the α2-adrenergic receptor mRNA in the ICo of male quail was examined.

## 2. Materials and methods

### 2.1. Animals

Twenty-seven male Japanese quail (*Coturnix japonica*) were obtained from the breeding colony at Azabu University. The ages were over 63 days post haching. All birds were sexually mature, as demonstrated by an enlarged cloacal gland and weighed from 90 to 150 g. They were maintained on a long-day photoperiod (16L:8D light/dark cycle; lights on 06:00) and provided with food and water *ad libitum*. They were housed in one room in individual wire cages (30 × 40 × 24 cm. Each bird was given a colored leg band for identification and randomly assigned to experimental groups. All birds had visual and auditory contact with other birds of the same sex. All experiments were approved by the Ethics Committee for the Use of Animals of Azabu University, Japan and follow ARRIVE guidelines (https://arriveguidelines.org/arrive-guidelines).

### 2.2. Stereotaxic surgery

Birds were anesthetized with an intramuscular injection of a ketamine (12.5 mg/kg body weight) and xylazine (25 mg/kg) mixture, and once sedated, placed in a stereotaxic apparatus (Narishige, Tokyo, Japan). Intracerebroventricular (ICV) cannulation was performed according to a stereotaxic atlas of the Japanese quail brain [22]. A stainless steel cannula guide sleeve (11 mm, 26 gauge; Plastics One, Akron, OH, USA) was inserted into the third ventricle (3.0 mm anterior, 0 mm lateral from the Y-point, and 7.0 mm deep from the surface of the dura mater). The cannula was anchored to the skull with resin cement (RelyX Unicem 2 Clicker; 3M ESPE, St. Paul, MN, USA). The cannula was kept closed at all times by a dummy wire affixed to a threaded cap. Correct placement of the cannula was confirmed 1 day after surgery by infusing 1 μg of human angiotensin-II (Calbiochem, San Diego, CA, USA) dissolved in 0.9% NaCl. This peptide rapidly stimulates drinking behavior in birds when infused into the third ventricle [23]. Only birds that immediately responded to the angiotensin-II infusion by increasing their water intake were used in the experiment.

### 2.3 Infusion and behavioral analysis

Using a within-subjects design, the effects of noradrenaline and clonidine on crow vocalizations were quantified in 18 Japanese quail, with 1 day between the tests. As the frequency of voluntary crowing during isolation differed among the birds (Fig. 1) the differences in the number of crows emitted by the same individual between days 1 and 2 of the behavioral experiment were examined, to determine the effects of the drugs on crow vocalizations. Birds were randomly assigned to one of six groups that received a central injection of 0.9% NaCl (vehicle), 1 or 10 μg of noradrenaline (norepinephrine bitartrate salt; Sigma-Aldrich, St. Louis, MO, USA), or 100 ng, 500 ng or 1 μg of clonidine (clonidine hydrochloride, Sigma-Aldrich) dissolved in vehicle. The birds were kept in an experimental cage (45 × 45 × 44.5 cm) in a soundproof box (MC-050T; Muromachi Kikai Co., Ltd., Tokyo, Japan) under 16L:8D (lights on 06:00) for at least 6 days before the start of the behavioral experiment. Food and water were provided to all birds *ad libitum* during the acclimation period and 60-min test periods. On day 1 of the experiment, the birds received infusions of the vehicle alone. The birds were gently held while a dummy wire was removed and the injector cannula, with the end of the tubing opposite to a Hamilton microsyringe, was inserted into the guide sleeve. Each bird was then returned to its experimental cage 1 min after the injection, and quail behavior was recorded by a Logicool HD Pro C920 webcam connected to a Windows PC via USB in addition to a Roland R‐26 portable recorder (Roland Corporation; 26-bits, sampling frequency: 44.1 kHz) during a 60-min period. On day 2 the birds received infusions of 1 μg of noradrenaline (n = 3), 10 μg of noradrenaline (n = 4), or 100 ng (n = 2), 500 ng (n = 2), or 1 μg of clonidine (n = 3) dissolved in vehicle or vehicle alone (n = 4) at the same time each bird received infusion of vehicle on day 1. After the injection, each bird was returned to its experimental cage. All solutions were injected over 10–20 s in a volume of 2 μl. All behavioral recordings were conducted between 09:00 and 12:00 to avoid any variability caused by diurnal rhythms in vocal behavior. The number of crow vocalizations was counted and the duration of food pecking was quantified. Three birds were selected from each of the vehicle and 1 μg clonidine groups for *in situ* hybridization and histological analyses based on their behavioral performance.

**Figure 1.**
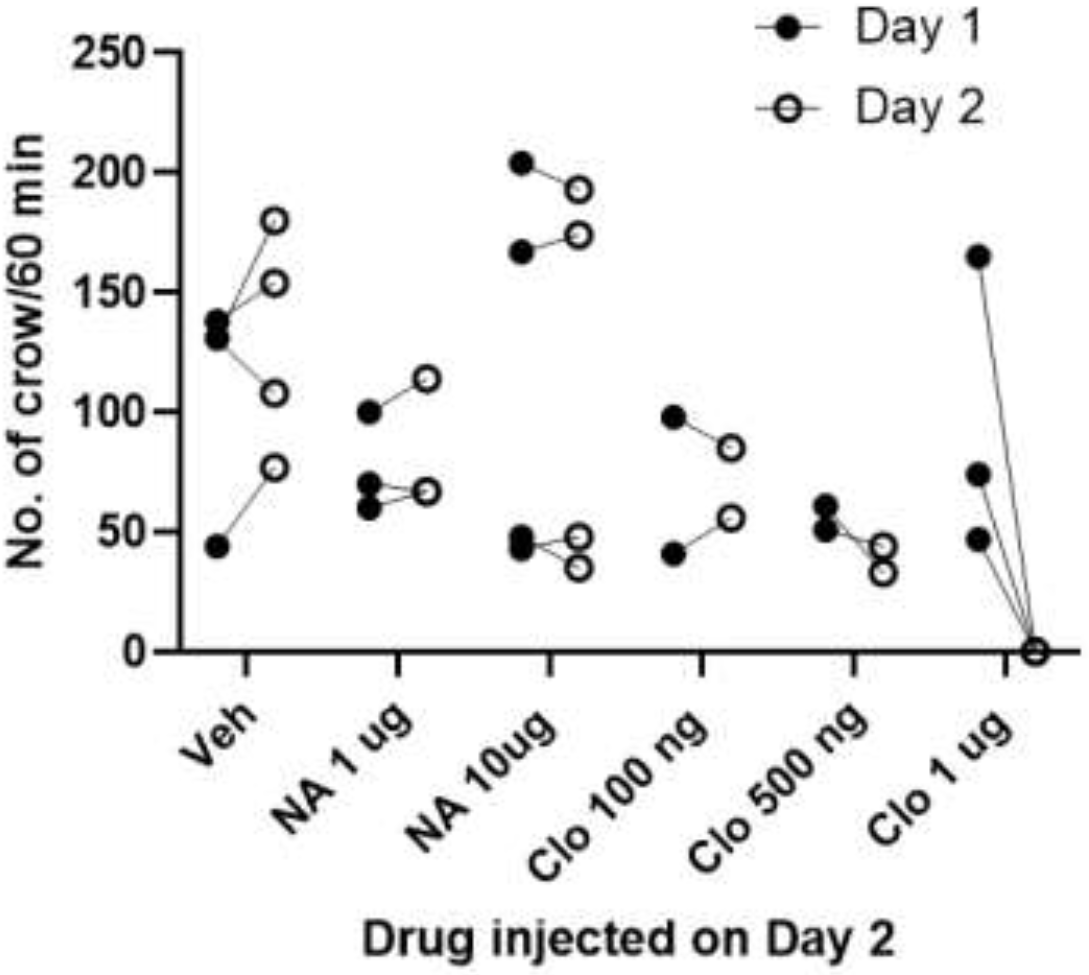
Frequency of male crowing during a 60-min test period after intracerebroventricular (ICV) injection of vehicle (Veh), noradrenaline (NA), or clonidine (Clo). Closed circles indicate the number of crow vocalizations performed by quail receiving an ICV injection of vehicle alone on day 1. Open circles indicate the number of crow vocalizations performed by quail receiving an ICV injection of vehicle, noradrenaline, or clonidine on day 2. Values for the same individual are connected by lines.

### 2.4. Immunohistochemistry

Three adult male quail in breeding condition were deeply anesthetized with mixed intramuscular injection of a ketamine (12.5 mg/kg body weight) and xylazine (25 mg/kg), and transcardially perfused with 0.01 M phosphate-buffered saline (PBS) followed by 4% paraformaldehyde. Brains were immediately dissected out of the skull and post-fixed in 4% paraformaldehyde in 0.1 M phosphate buffer, pH 7.4 for 1 day at 4°C and then transferred into 30% sucrose in 0.1 M phosphate buffer, pH 7.4 solution and kept for 3 days at 4°C. Brains were then embedded in Tissue-Tek optimal cutting temperature compound (Sakura Finetek, Tokyo, Japan), frozen on powdered dry ice and stored at −80°C until sectioning. Each brain was then sectioned in the frontal plane at 50 μm on a cryostat (Leica Microsystems, Wetzlar, Germany), and every sixth section through the brainstem from each animal was collected into 0.1M PBS. Free-floating sections were rinsed for 5 min in PBS and then incubated for 15 min in PBS containing 0.6% hydrogen peroxide to inhibit endogenous peroxide activity. Next, the sections were rinsed three times for 5 min with 0.3% Triton X-100 in PBS (PBST) and blocked with 5% normal sheep serum in PBST for 2 h at room temperature (RT). The sections were then incubated overnight with rabbit polyclonal antiserum against dopamine-beta-hydroxylase (DBH) purified from the bovine adrenal medulla (Product ID: 22806; ImmunoStar, Hudson, WI, USA) at a dilution of 1:2,000 in PBST and 4°C. DBH is a specific marker for noradrenergic neurons in the brain. The specificity of the antiserum has been described previously [24]. After rinsing in PBST three times for 5 min, the sections were incubated in biotinylated goat anti-rabbit IgG (H+L) ((Product ID: BA-1000; Vector Laboratories, Burlingame, CA, USA) in PBST at a dilution of 1:1,000 for 2 h at RT. The sections were then washed with PBS three times for 5 min and incubated with avidin-biotin complex reagent at a dilution based on the manufacturer’s recommendations (Vector Laboratories) for 30 min at RT. After washing in PBS three times for 5 min and rinsing with 0.1 M Tris-HCl buffer (pH 7.6), the sections were stained with 0.02% 3,3-diaminobenzidine substrate solution (Dojindo Molecular Technologies, Kumamoto, Japan) for 5 min. After washing in PBS three times for 5 min, the sections were mounted on 3-aminopropyltriethoxysilane-coated slides (Matsunami Glass, Osaka, Japan) and dried overnight at RT. The slide-mounted sections were then dehydrated through a graded ethanol series and coverslipped with Entellan mountant (Merck, Darmstadt, Germany). All sections were viewed under a bright field microscope and images of sections were taken with a Keyence BZ-X710 microscope (Keyence Corp., Osaka, Japan).

### 2.5. In situ hybridization

The cDNA fragments [α2A-adrenergic receptor (GenBank Acc. No. AB820133), α2C-adrenergic receptor (GenBank Acc. No. AB820134), and zenk (GenBank Acc. No LC622525)] were isolated from the adult midbrain of two adult male Japanese quail in breeding condition by reverse transcription-polymerase chain reaction. The following primers were used: [5′-CAACGTCCTGGTCATCATTG-3′ and 5′-ATGACGAAGACCCCAATCAC-3′ for α2A-adrenergic receptor, 5′-CTCTGGTCATGCCTTTCTCC-3′ and 5′-TCCCGGCAAATACCATAGAG-3′ for α2C-adrenergic receptor; and 5′-AAAACCATGCCAGAAACCAG-3′ and 5′-GGCAGCAACAGAGGAAGAAG-3′ for zenk. Each cDNA fragment was inserted into the pCR II-T (Invitrogen, Carlsbad, CA, USA) or p-GEM-t easy vectors (Promega, Madison, WI, USA). The plasmids were digested with the XhoI, SpeI, or NcoI enzymes to release the fragment, and probes were synthesized using SP6 or T7 RNA polymerase (Roche Diagnostics, Rotkreuz, Switzerland) with a digoxigenin (DIG)-labeling mix (Roche Diagnostics). Sense probes corresponding to each antisense probe were also synthesized as controls.

Four adult male quail in breeding condition were killed by rapid decapitation for analyses of α2A- and α2C-adrenergic receptor mRNA expression. Six quail (vehicle-injected quail, n = 3; clonidine-injected quail, n = 3) were immediately decapitated (within 5 min after the end of the behavioral test) to quantify zenk mRNA expression in the ICo. Each brain was carefully removed from the skull and embedded in Tissue-Tek optimum cutting temperature compound (Sakura Finetek), frozen on dry ice and stored at –80°C until analyses. Frozen brains were cut into 20-μm coronal sections on a cryostat (Leica Microsystems). Every sixth section through the midbrain from each animal was mounted on 3-aminopropyltriethoxysilane-coated slides (Matsunami Glass) and stored at –80°C until use. The sections were post-fixed in 4% paraformaldehyde for 10 min and washed in 0.1 M PBS (pH 7.4) three times for 5 min. The slides were then acetylated with 0.25% acetic anhydride in 0.1 M triethanolamine for 10 min and washed in PBS with 1% Triton X-100 (Sigma-Aldrich) for 30 min. The sections were then incubated at RT with hybridization buffer containing 50% formamide (Wako Pure Chemicals, Osaka, Japan), 5× saline sodium citrate, 1× Denhardt’s solution (Sigma-Aldrich), 200 μg/mL yeast tRNA (Roche Diagnostics), and 500 μg/mL DNA (Roche Diagnostics). The sections were hybridized at 72°C overnight in a hybridization buffer with RNA probes. The sections were rinsed in 0.2× saline sodium citrate at 72°C for 2 h and then blocked for 2 h in a solution of 0.1 M Tris (pH 7.5, Pure Chemicals) and 0.15 M NaCl (Pure Chemicals) with 10% sheep serum. The slides were incubated overnight with a 1:5,000 dilution of alkaline phosphatase-conjugated anti-DIG antibody (Roche Diagnostics). After the slides had been washed in in a solution of 0.1 M Tris (pH 7.5) and 0.15 M NaCl with 0.1% Polysorbate 20 (Pure Chemicals) three times for 5 min, alkaline phosphatase activity was detected by incubation with 375 mg/mL nitroblue tetrazolium chloride (Roche Diagnostics) and 188 mg/mL 5-bromo-4-chloro-3-indolyl phosphate (Roche Diagnostics) in a solution of 0.1 M Tris (pH 9.5), 0.1 M NaCl, and 50 mM MgCl_2_ (Pure Chemicals) at RT overnight. This results in labeled cells expressing a purple-blue color. Brain sections incubated with control sense probes processed in the same way as detailed above showed no specific reactivity. Images of ICo sections were captured with a Keyence BZ-X710 microscope or a LeicaMC170 HD camera attached to a Leica DM500 microscope (Leica Microsystems).

### 2.6 Statistical analyses

Statistical analyses were conducted with Prism 8.0 software (GraphPad Software, La Jolla, CA, USA) or IBM SPSS Statistics v24 (IBM Corp., Armonk, NY, USA). Data were analyzed by a one-way analysis of variance (ANOVA) followed by Tukey’s multiple comparison test or a repeated two-way ANOVA followed by Bonferrini’s test. *P* values < 0.05 were considered significant.

## 3. Results

### 3.1. Effects of noradrenaline and clonidine on crow vocalizations

ICV injection of noradrenaline had no effect on male crowing behavior at either dose during the 60-min test period compared to vehicle treated birds (Figs. 1 & 2). In contrast, 0.5 µg of clonidine inhibited crow vocalizations 20 min after injection, and 1 µg of clonidine had the same effect for 60 min (Fig. 3). Clonidine decreased the frequency of crowing in a dose dependent manner (F_(5,12)_ = 5.466, *P* = 0.0075 by one-way ANOVA, *P* = 0.0043 vs. vehicle, *P* = 0.047 vs. 100 ng of clonidine by Tukey’s multiple comparison test; Fig. 2).

**Figure 2.**
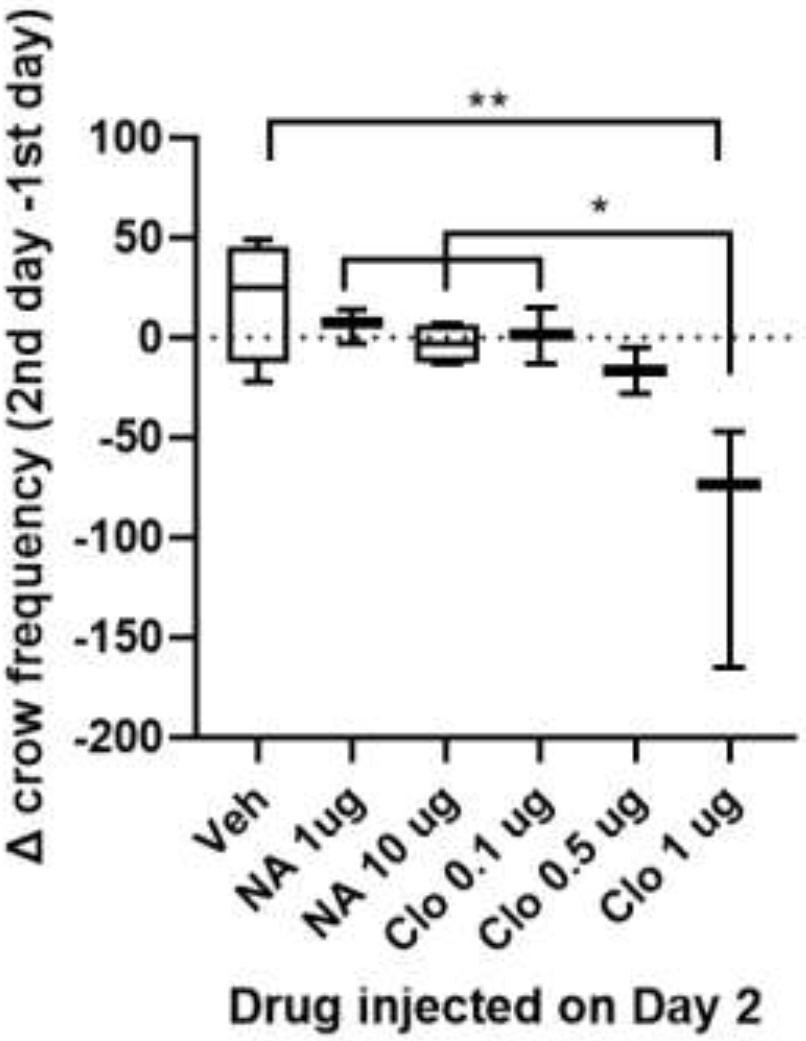
Effects of clonidine on the frequency of crowing. Clonidine-injected quail decreased the number of crows on day 2 in a dose-dependent manner compared with the control on day 1. Box plots show the median (line) and 75^th^ and 25th percentiles (box) with the 95% confidence interval (error bars). NA, noradrenaline; Clo, clonidine; Veh, vehicle. * *P* < 0.05, ** *P* < 0.01.

**Figure 3.**
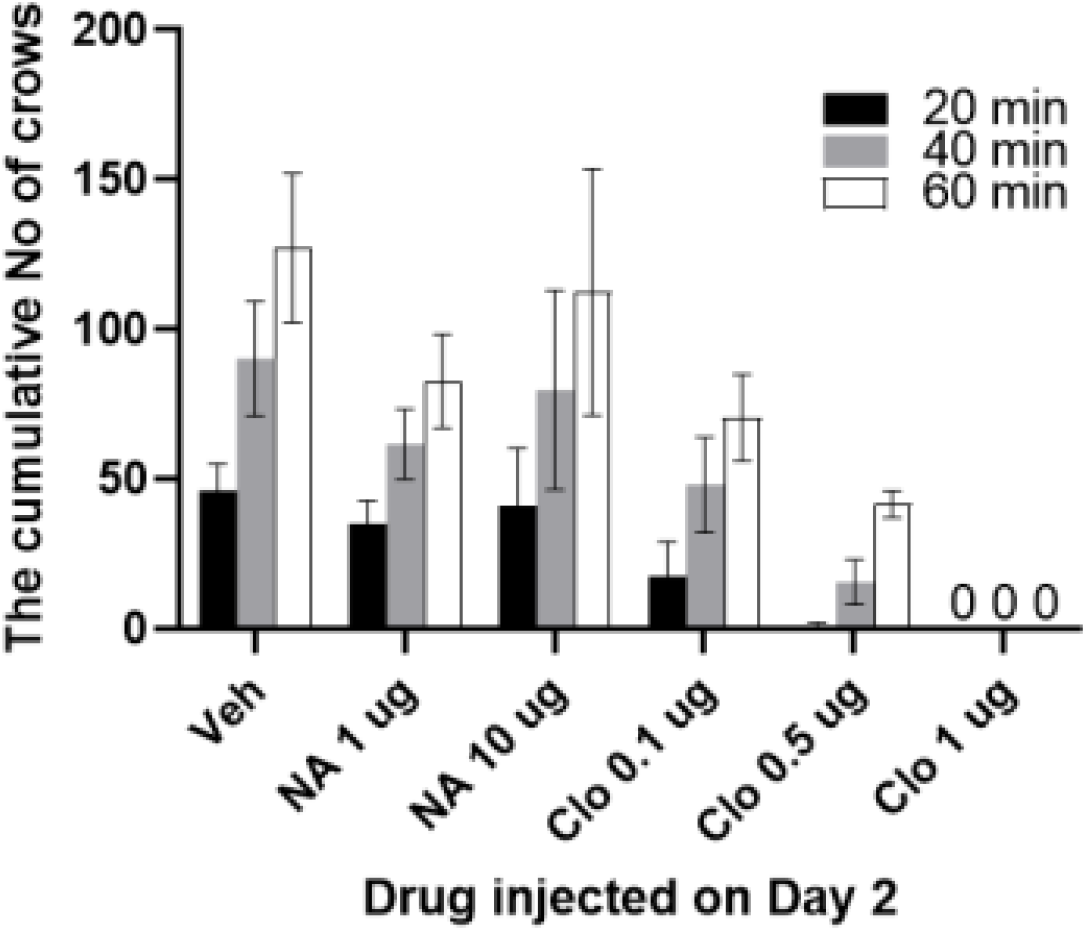
Cumulative number of crows over a 1 h-period after an intracerebroventricular (ICV) injection of vehicle, noradrenaline or clonidine in male quail. Values are means ± SEM. NA, noradrenaline; Clo, clonidine; Veh, vehicle.

Measures of feeding behavior were analyzed via a two-way repeated measures ANOVA with group (clonidine/vehicle) and day (Day 1/Day 2) as factors and central administration of 1 μg of clonidine increased feeding behavior compared to vehicle administration. (Fig. 4). There was a main effect in the group factor (F_(1, 5)_ = 10.10, *P* = 0.025) and in the day factor (F_(1, 5)_ = 13.00, *P* = 0.015). In the interaction analysis between group and day factors, a significant interaction was found (F_(1, 5)_ = 12.76, *P* = 0.016). A simple main effect test using the Bonferroni method revealed significant differences between the vehicle - injected birds on Day 1 and the clonidine-injected birds on Day 2 (within subject; *P* = 0.005) and between the vehicle and clonidine-injected on Day 2 (between subject; *P* = 0.019).

**Figure 4.**
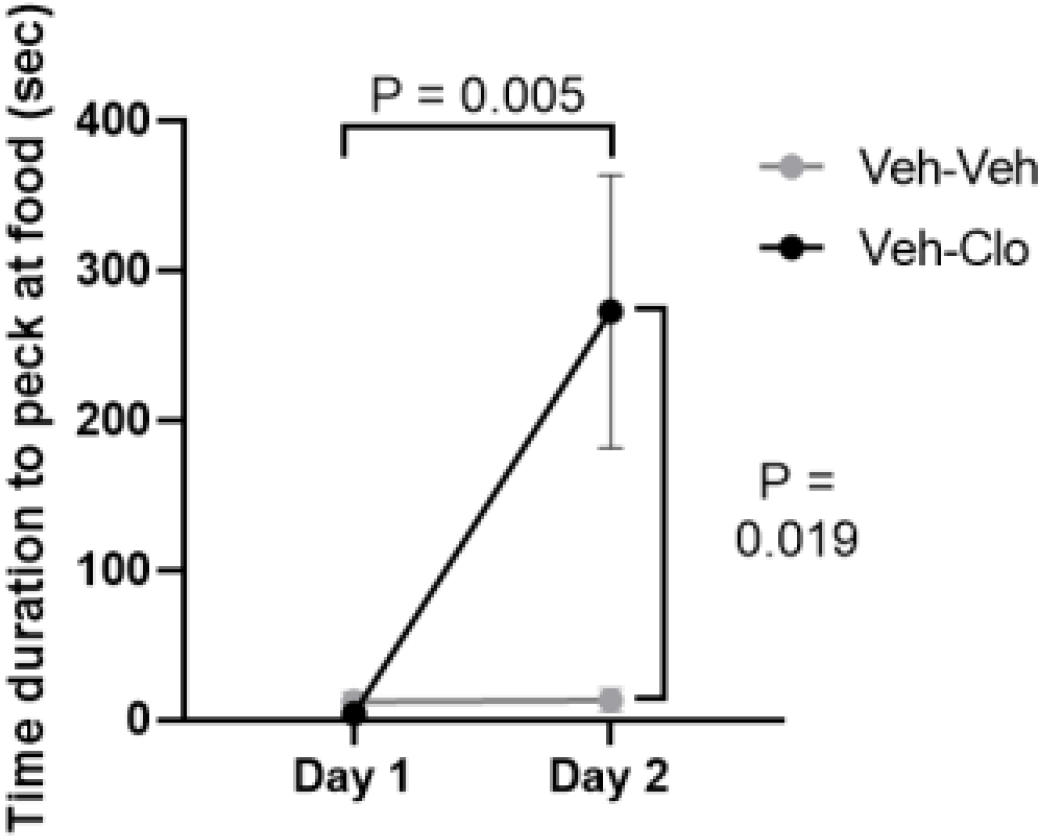
Effect of clonidine on feeding behavior. Duration of pecking of food by quail that received an intracerebroventricular (ICV) injection of either vehicle (gray circles) or clonidine (black circles) on days 1 and 2. Values are means ± SEM and for the same group are connected by lines. NA, noradrenaline; Clo, clonidine.

### 3.2. Zenk mRNA expression after ICV clonidine injection

Zenk mRNA expression was detected in the dorsomedial ICo of all the ICV-vehicle injected quail (Fig. 5A) but 1 µg ICV-clonidine completely abolished this expression (Fig. 5B). No signal above background was observed using the zenk sense control probes (Fig. 5C & D).

**Figure 5.**
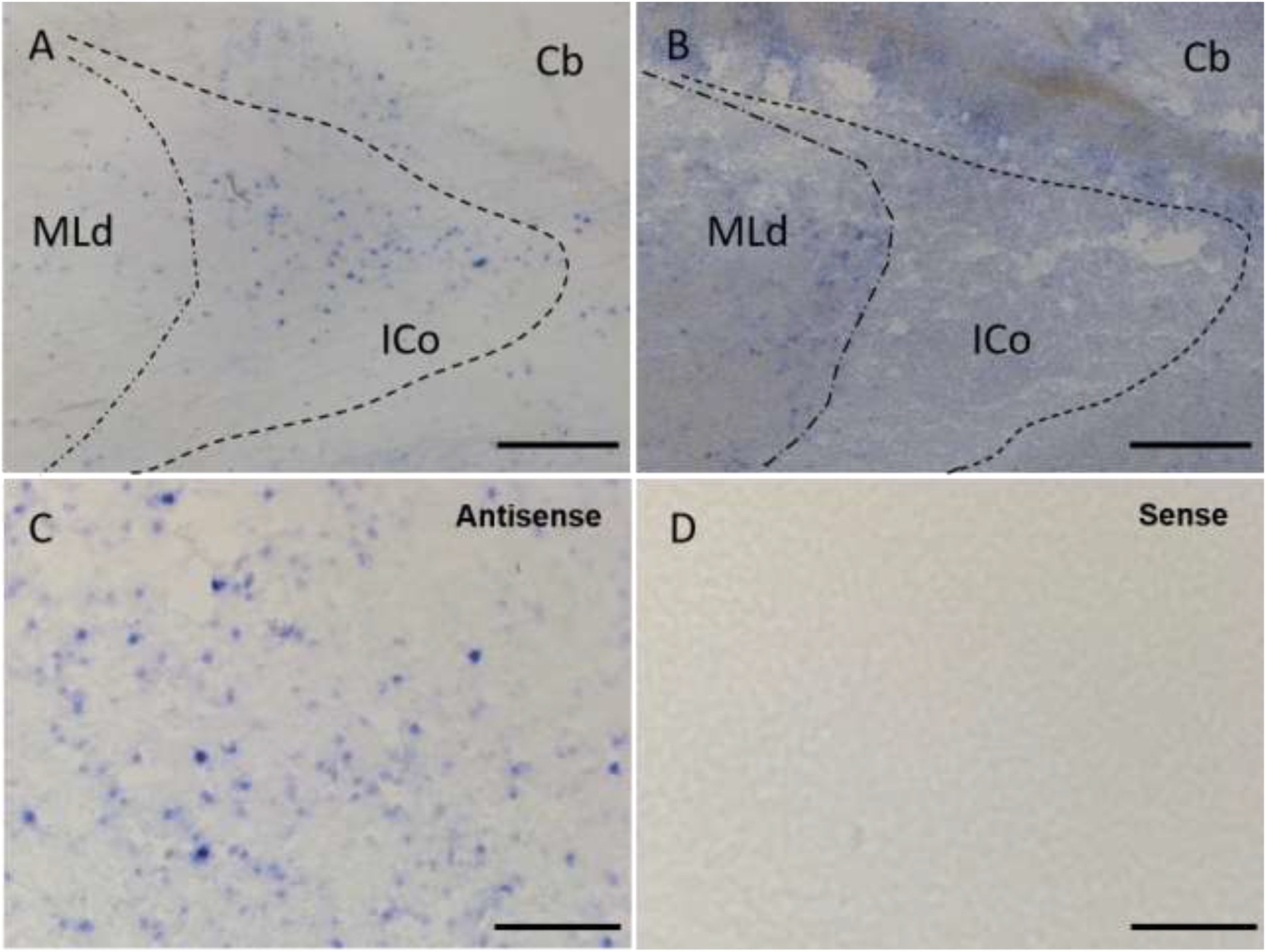
Effect of clonidine on zenk mRNA expression in the midbrain vocal center. Representative photomicrographs depicting the effect of vehicle (A) and clonidine (B) on zenk mRNA expression. Hybridization with sense zenk probes served as a negative control for antisense probe specificity in vehicle ICV injected birds (C, D). Scale bars = 500 μm (A & B), 200 μm (C & D). Cb, cerebellum; ICo, nucleus intercollicularis; MLd, nucleus mesencephalicus lateralis pars dorsalis.

### 3.3. Noradrenergic innervation of the dorsomedial ICo

We have previously used DBH immunostaining as a noradrenergic marker and described antiserum specificity [24]. The same immunostaining method was applied this study. DBH-immunoreactive cell bodies were observed in the locus coeruleus (LoC; Fig. 6A), nucleus subcoeruleus ventralis (SCv; Fig. 6B), and lateral tegmental field (LT; Fig. 6C), as well the DBH-immunoreactive fibers in the dorsomedial ICo (Fig. 6D), which possessed many varicosities (Fig. 6E).

**Figure 6.**
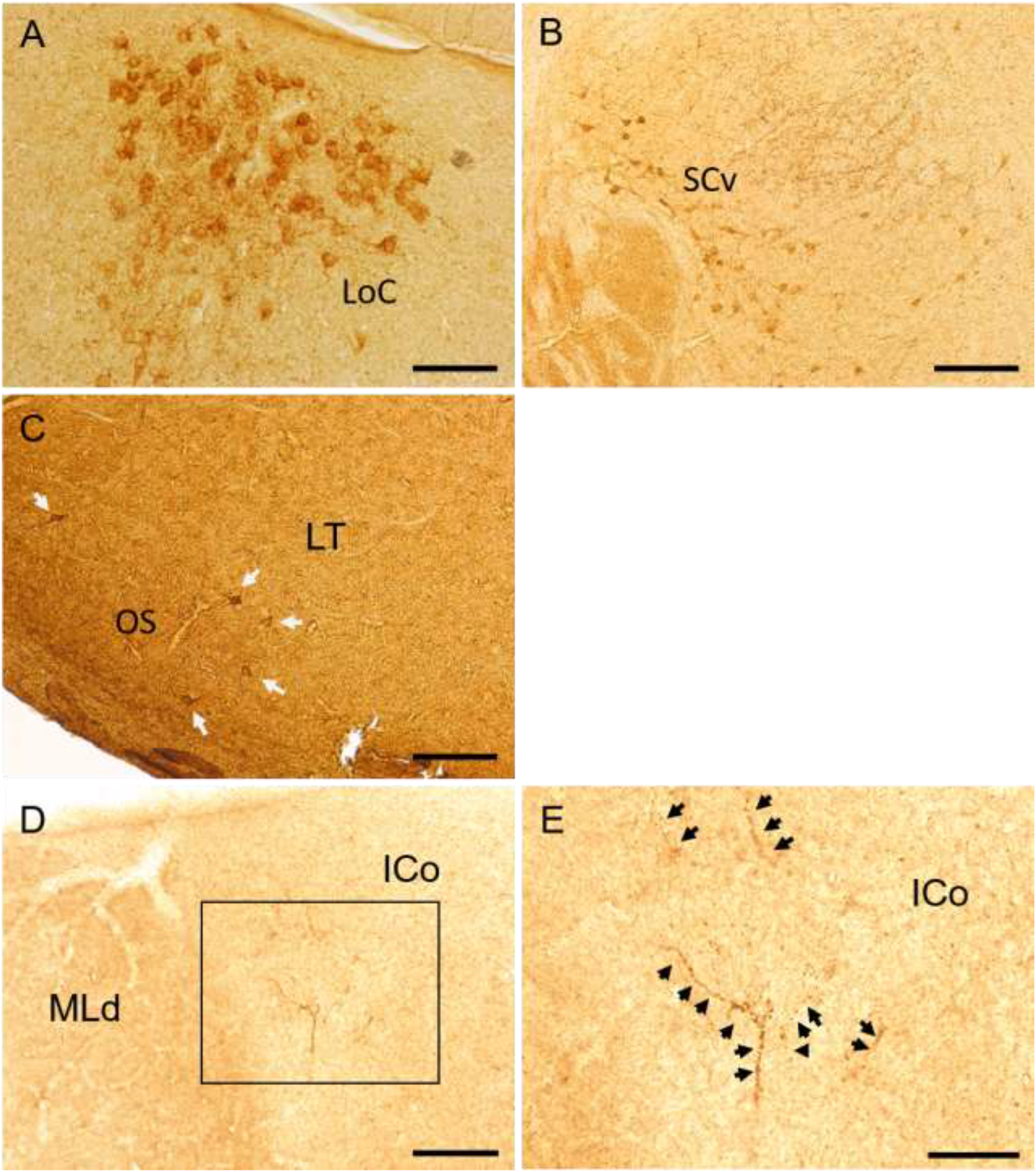
Photomicrographs of dopamine beta hydroxylase (DBH)-immunoreactive (ir) cells and fibers. DBH-ir cells in the locus coeruleus (LoC, A), nucleus subcoeruleus ventralis (SCv, B), and lateral tegmental field (LT, C). Low-power photomicrograph of the DBH-ir fibers in the nucleus intercollicularis (ICo, D). Higher magnification photomicrograph of the DBH-ir fibers in the dorsomedial ICo (E). Scale bars = 100 μm (A & D), 200 μm (B & C) and 500 μm (E).

### 3.4. Expression of α2-adrenergic receptor mRNA in ICo neurons

*In situ* hybridization identified α2-adrenergic receptor mRNA expression in the ICo of the Japanese quail. cDNAs from the midbrain were cloned for α2A- and α2C-, but not α2B-adrenergic receptor, and as templates to prepare *in situ* hybridization probes. Intense expression of α2A-adrenergic receptor mRNA was localized in the dorsomedial ICo (Fig. 7A). Signal for α2C-adrenergic receptor mRNA expression were found in the optic tectum, but not in the ICo (Fig. 7C). Controls, in which the sense RNA probes were substituted for the antisense RNA probes, showed no positive mRNA signal in any of the brain sections (Fig. 7B & D).

**Figure 7.**
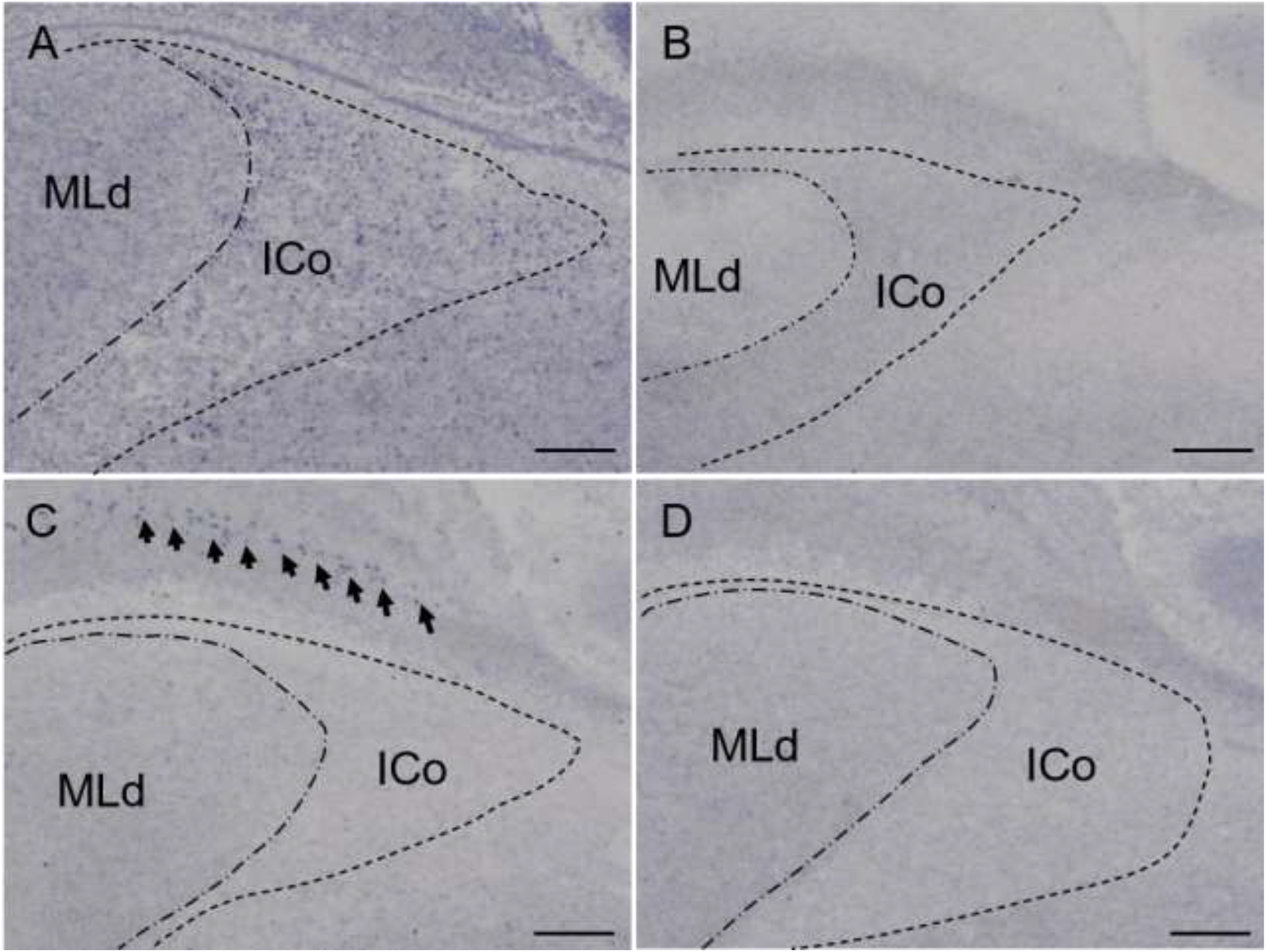
α2A and α2C-adrenergic receptor mRNA expression in the midbrain. Representative α2A- and α2C-adrenergic receptor mRNA expression by *in situ* hybridization (A, C). Hybridization with sense α2A- and α2C-adrenergic receptor probes served as a negative control for antisense probe specificity (B, D). Scale bars = 500 μm.

## 4. Discussion

Centrally administered noradrenaline decreases plasma luteinizing hormone concentrations in adult male quail in breeding condition [23], which in turn possibly reduces blood testosterone concentrations. Crowing in quail is androgen-dependent [25], and we expected to observe an inhibitory effect of noradrenaline on crowing *via* activation of the hypothalamus-pituitary-gonad axis as well as a direct effect on the α2-adrenergic receptors expressing in the ICo but those were not observed. Our results demonstrate for the first time that ICV-administered clonidine, but not noradrenaline, powerfully suppressed crowing in male adult quail in a dose-dependent manner. This finding may be explained by the fact that clonidine diffuses into the ventricles and binds to α2-adrenergic receptors, more potently than noradrenaline.

Clonidine has sedative effects, which is mediated by activation of central α2-adrenergic receptors but central administration of clonidine does not reduce all behavioral activity. In the present study, ICV injection of clonidine stimulated feeding behavior of adult male quail and this effect was similar to previous studies conducted using broiler and layer-type chick [26, 27]. An α2-adrenergic receptor antagonist, yohimbine attenuated the clonidine induced food intake [27], indicating that α2-adrenergic receptors are related to the stimulation of feeding in chick. Since stimulation of food intake caused by neuropeptide Y and beta-endorphin was attenuated by co-injection with yohimbine, the receptors also likely mediate the orexigenic effects of neuropeptide Y- and beta-endorphin in layer-type chicks [27]. These suggest that NPY- or endorphin-producing neurons form neural circuits that control food intake with noradrenergic neurons acting on α2-adrenergic receptors in the Galliformes species.

Activation of neurohormone and neurotransmitter receptors initiate an intracellular signaling cascade that leads to changes in genes transcription. The IEGs are critical signaling intermediates within this cascade and modulate neuronal activities by controlling gene transcription, *via* the binding of their protein forms to regulatory sites on DNA [21, 28]. Noradrenaline has a documented role in the regulation of IEG expression as lesioning noradrenergic fibers with the noradrenergic neurotoxin N-(2-chloroethyl)-N-ethyl-2-bromobenzylamine results in changes in telencephalic and hypothalamic zenk expression in adult birds [29, 30]. The present study demonstrated that zenk mRNA was expressed in the dorsomedial region of the ICo in vehicle-injected males after they emitted crows. In comparison, male crowing was abolished by ICV-administered clonidine and this resulted in reduced zenk mRNA expression in the dorsomedial ICo. The neuronal expression of zenk mRNA in male quail, which emitted high-frequency crows, and their reduction by clonidine, suggests a link between the zenk gene in ICo neurons and the regulation of crowing. The reduction of zenk mRNA expression by clonidine may reflect a direct inhibitory action of clonidine on the dorsomedial ICo neurons through α2-adrenergic receptors to cause complete loss of crowing.

To investigate whether noradrenaline directly controls the midbrain vocal nucleus of ICo, the noradrenaline neuronal marker DBH was immunohistologically stained in the male quail brain. DBH-immunoreactive cell bodies were localized in the LoC, SCv, and LT, and DBH-immunoreactive fibers, were localized in the dorsomedial part of the ICo of male quail. A tract-tracing study in pigeons [31] showed that the ICo receives projections from LoC and SCv neurons. Together these findings suggest that ICo neurons receive noradrenergic innervation, mainly from the LoC and SCv. Only α2A-adrenergic receptor mRNA was expressed in the dorsomedial region of the ICo and α2C-adrenergic receptor mRNA was expressed in the optic tectum, which is part of the midbrain visual system. These histochemical results support a direct inhibitory effect of noradrenaline on ICo neurons through the α2A-adrenergic receptor.

Noradrenaline is thought to play an important role in the behavioral regulation of potential mate cues such as arousal, attention, and goal-directed approach responses [29, 32, 33]. Male quail crow frequently during the breeding season as crowing elicits phonotaxis in female quail [2] but crowing is suppressed in the presences of a female. When male quail view a female there is increased extracellular noradrenaline release in the brain [23]. Therefore, it may be hypothesized that the presence of the female rapidly increases extracellular noradrenaline release in the brain, which acts on ICo neurons *via* the α2A-adrenergic receptor to inhibit ICo neuronal activity and suppress crowing. However, further research is required to test this hypothesis.

## 5. Conclusion

Together these data suggest that the α2-adrenergic system in the midbrain vocal center suppresses courtship vocalizations in male quail as centrally administered clonidine suppressed male crowing in a dose-dependent manner and diminished zenk mRNA expression in the dorsomedial ICo. We conclude that noradrenaline is likely to be a major factor in the regulation of bird vocalizations as the ICo receives noradrenergic innervation and express α2A-adrenergic receptor mRNA.

## Funding

The present study was supported by Grants-in-Aid for Scientific Research from the Japan Society for the Promotion of Science (KAKENHI, JP16K18585) and Ministry of Education, Culture, Sports, Science and Technology-Supported Program for the Private University Research Branding Project to Y. Tobari. S. L. Meddle acknowledges Roslin Institute Strategic grant funding from the Biotechnology and Biological Sciences Research Council (BBSRC), UK (BB/P013759/1) and a BBSRC Japan Partnering Award (BB/M027805/1).

## CRediT authorship contribution statement

**Yasuko Tobari**: Conceptualization, Formal analysis, Validation, Writing-original draft, Visualization, Project administration, Funding acquisition. **Ami Masuzawa**: Formal analysis, Investigation, Visualization. Validation.**Norika Harada**: Investigation, Visualization. **Kenta Suzuki**: Formal analysis. **Simone L Meddle**:Writing-original draft, Funding acquisition.

## Declaration of Competing Interest

The authors declare no conflicts of interest.

## Acknowledgements

We thank Mr. Yukinori Ebihara and Ms. Chie Ohya for their technical assistance.

## Notes

### Competing Interest Statement

The authors have declared no competing interest.

